# Effects of animal dormancy on oxidative stress, immune status, and glucocorticoids: a meta-analysis

**DOI:** 10.1101/2025.10.02.680078

**Authors:** Pablo Burraco, Tamara G. Petrović, Pablo Capilla-Lasheras, Marko Prokić

**Affiliations:** Department of Ecology and Evolution, Doñana Biological Station, Spanish National Research Council (CSIC). Avenida Americo Vespucio 26, 41020 Sevilla (Spain); Department of Physiology, Institute for Biological Research Siniša Stanković, National Institute of the Republic of Serbia, University of Belgrade. Bulevar Despota Stefana 142, Belgrade, 11060, Serbia; Bird Migration Unit, Swiss Ornithological Institute, Sempach, Switzerland; School of Biodiversity, One Health and Veterinary Medicine, University of Glasgow, Glasgow, United Kingdom

**Keywords:** adaptive immunity, aestivation, arousal, hibernation, hypoxia, innate immunity, metabolic depression, oxidative stress, thermoregulatory strategy, torpor

## Abstract

Despite the growing interest of evolutionary biologists and ecologists in unravelling the mechanisms behind animal responses to challenging conditions, our knowledge remains limited. To contribute to this field, we conducted a global meta-analysis (N= 112 unique studies, k = 2,840 independent estimates) to synthesize empirical information on the influence of vertebrate dormancy (*i.e.,* cold- and heat-induced dormancy) on redox status, immune function, and glucocorticoid levels. Our study reveals that dormancy has contrasting effects on animal physiology, depending on the thermoregulatory strategy. In endotherms, both dormant and arousal states are accompanied by reduced ROS production, lower oxidative damage, or upregulated antioxidant response, whereas in ectotherms, greater oxidative damage occurs only during the dormant state. Likewise, innate immune responses are upregulated during arousal in endotherms, while ectotherms downregulate pathways involved both in innate and adaptive responses during the dormant state. Interestingly, glucocorticoid levels remain stable throughout dormancy, suggesting that undergoing metabolic shifts may uncouple glucocorticoid release. These findings provide mechanistic explanation on the eco-evolutionary consequences of dormancy, revealing diverging strategies between endotherms and ectotherms. By highlighting metabolic depression as a central axis of these responses, our results also inform conservation strategies for dormant species facing rapid environmental change.

## Introduction

Organisms have evolved very diverse strategies to cope with challenging environments. In this context, animal dormancy represents a fascinating adaptation to survive prolonged periods of adverse conditions such as extreme temperatures (Geiser, 2013; Jiang *et al*., 2023). In response to cold conditions, many endotherms enter hibernation (*i.e.,* prolonged bouts of torpor), while torpor (*i.e.,* short-term, reversible reductions in body temperature and metabolism) occurs in small endotherms. Additionally, many ectotherms, along with some endotherms, undergo aestivation in response to extreme heat and drought. The onset of dormancy includes abrupt reductions in organismal activity and metabolic rates, which in some taxa may be associated with hypoxia or anoxia (Storey and Storey, 1990; Staples, 2016). Such drastic changes are thought to broadly affect animal physiology. In endotherms, arousal leads to the progressive return to euthermia and normal activity, whereas ectotherms resume activity as environmental conditions improve. To what extent dormancy perturbs or reshapes organismal physiology is still under debate. Comparative syntheses across taxa provide critical insights into the eco-evolutionary relevance of dormancy and guide future work aiming to unravel its mechanistic basis, while also improving predictions of how dormant species will respond to rapid climate change.

Organisms entering the dormant state experience dramatically reduced oxygen availability for extended periods, a condition that is largely accommodated by a marked reduction in metabolic activity, known as metabolic depression (Mohr, Bagriantsev and Gracheva, 2020; Staples, Mathers and Duffy, 2022; Jiang *et al*., 2023). Lowered metabolism reduces mitochondrial respiration (Ali *et al*., 2010; Salin *et al*., 2016) which would be expected to decrease the production of reactive oxygen species (ROS) (Storey and Storey, 1990). However, empirical studies have reported elevated ROS levels in some hypoxic, dormant organisms (Brown *et al*., 2012; Yin *et al*., 2016; Wei *et al*., 2018). This apparent paradox suggests that alternative sources of ROS may remain active during dormancy or that transient ROS production serves as a signal to trigger antioxidant defences. Indeed, transcription factors that upregulate antioxidant capacity are often activated in dormant organisms (Carey, Frank and Seifert, 2000; Wu and Storey, 2016; Staples, Mathers and Duffy, 2022). In line with this, the *Preparation for oxidative stress* (POS) theory proposes that enhanced antioxidant defences during dormancy prepare organisms to counteract oxidative stress that arises upon arousal, when the transition from anoxic (dormant) to normoxic (post-dormancy) conditions occurs (Hermes-Lima *et al*., 2015; Moreira *et al*., 2016; Giraudeau *et al*., 2018). It remains uncertain whether POS during hypoxia is universal, and whether antioxidant action alone is sufficient to neutralise ROS and prevent oxidative damage throughout dormancy, including both the dormant and arousal states (Moreira *et al*., 2016; Giraud-Billoud *et al*., 2024). While enhancing antioxidant defences is generally considered an energetically costly process, the observation that some studies report minimal cell death or organ injury in dormant individuals suggests that evolution may have minimized these costs to mitigate oxidative stress (Bouma *et al*., 2012).

Immune function is a key determinant of animal life histories (McDade, 2003; Cooper and Alder, 2006) and is thought to be influenced by dormancy (Bouma *et al*., 2012). Changes in the immune status can take place either via the release of ROS that can act as signalling molecules triggering immune responses, or through shifts in energy allocation away from immune maintenance towards other physiological pathways (Bouma *et al*., 2012). In some organisms, dormancy result in inflammation, then leading to tissue dysfunction and injury (Cain and Cidlowski, 2017). Suppression of immune activity (both innate and adaptive) has been also found during dormancy states (Mohr, Bagriantsev and Gracheva, 2020; Jiang *et al*., 2023). For example, dormant animals can experience a reduction in circulating leukocytes up to 90%, as well as decreased phagocytic activity or antibody production (Bouma *et al*., 2012). Upon arousal, the expression of several immune pathways returns to baseline levels in some species (Bouma *et al*., 2012). Whether these immune responses are universal in dormant organisms is unknown despite this can help to understand dormancy impacts on animal health and fitness.

As with the putative effects of dormancy on redox status or immune function, the release rate of certain hormones, particularly those involved in the regulation of metabolism, may also be highly sensitive. In vertebrates, high levels of their main glucocorticoids (cortisol in mammals and fish, and corticosterone in birds, reptiles, and amphibians) are found in animals coping with detrimental environmental conditions which can shape life histories (Crespi *et al*., 2013; Romero and Beattie, 2022). Glucocorticoid release can also enhance metabolism and result in an oxidative stress state and immunosuppression (Vegiopoulos and Herzig, 2007; Denver, 2009; Costantini, Marasco and Møller, 2011). In dormant vertebrates, whether glucocorticoid levels are higher such as often observed in stressed individuals, or lower hence aligning with metabolism depression, requires a global evaluation regarding the diversity of studies conducted to date.

Dormancy effects on animal physiology may be shaped by the thermoregulatory capacity of organisms. Endotherms may cope more effectively with dormancy than ectotherms, whose capacity to adjust internal body temperature is limited. Furthermore, the effects of dormancy on redox balance may be influenced by an animal’s thermoregulatory strategy. Endotherms are likely to cope more effectively with dormancy than ectotherms, whose ability to regulate internal body temperature is limited. Investigating physiological changes (and particularly oxidative stress, immune responses, and glucocorticoids) across taxa undergoing dormancy is therefore critical for understanding their consequences on animal life history, such as reproductive success and lifespan (Monaghan, Metcalfe and Torres, 2009; Metcalfe and Alonso-Alvarez, 2010; Costantini, 2024).

We conducted a systematic review and meta-analysis to assess variation in redox status (ROS, regulatory pathways, antioxidants, and markers of oxidative damage), immune function (innate, adaptive, or pathways involved in both), and glucocorticoid levels (cortisol and corticosterone) during dormant and arousal states in hibernating or aestivating vertebrates. We hypothesize that dormancy and arousal will alter redox balance, with the direction and magnitude of oxidative stress and antioxidant responses differing between endotherms and ectotherms, as ectotherms may experience greater oxidative stress due to their limited thermoregulatory capacity. Similarly, immune function is expected to be modulated during dormancy, with endotherms potentially maintaining or even upregulating certain pathways during arousal, whereas ectotherms may downregulate both innate and adaptive responses. Glucocorticoid levels (cortisol or corticosterone) are anticipated to remain largely stable across dormancy, with possible transient increases during arousal. By synthesizing these patterns across taxa, this study aims to identify general predictors of physiological changes during dormancy, offering insights with eco-evolutionary, biomedical, and conservation relevance.

## Methods

### Literature review and criteria for inclusion

We carried out a systematic literature search to collect available empirical data on the effect of dormancy states on animal oxidative stress, immune responses and glucocorticoid levels. For oxidative stress, we used the search string ‘oxidative stress’ AND ‘hibernation’ OR ‘torpor’ OR ‘estivation’ OR ‘aestivation’. For immune system, we used the search string ‘immune’ AND ‘hibernation’ OR ‘torpor’ OR ‘estivation’ OR ‘aestivation’. For glucocorticoids, we used the search string ‘cortisol’ OR ‘corticosterone’ AND ‘hibernation’ OR ‘torpor’ OR ‘estivation’ OR ‘aestivation’. All the searches were performed in Web of Science, PubMed, and Scopus databases. Preferred Reporting Items for Systematic Reviews and Meta-Analyses (PRISMA) diagrams are included in Supplementary Material S1-S3.

Papers and data had to report the following information to be included in the meta-analyses: i) mean oxidative stress, immune, or glucocorticoid values, variation (standard deviation or standard error) and sample size, measured ii) at least across two dormant phases (*i.e.,* euthermia, hibernation, torpor, or estivation, or arousal), in iii) animals do not exposed to any other potentially stressful condition (*e.g.,* pollutants or predators). For oxidative stress, we collected data on several components of the antioxidant system: enzymatic and non-enzymatic antioxidants, reactive oxygen species, markers of lipid and protein oxidative damage, and expression of regulatory pathways. For immune function, we collected data of parameters of the adaptive, innate or both immune systems. For glucocorticoids, we collected, cortisol and corticosterone values. The full list of markers can be found in the available dataset (see data accessibility statement below). If data met the requirements to be included in the meta-analysis, we also collected the following information: year of the study, species name, thermoregulatory mechanism (*i.e.,* ectothermic or endothermic), location where the study was conducted, experimental venue (*i.e.,* field, laboratory or captivity), biological process (*i.e.,* hibernation or estivation), biological matrix (tissue in which the physiological markers were measured), name of the specific marker measured, developmental stage, the unit of the measured marker, and the timing of the state (‘early’ if animals spent five or less days within that state, otherwise ‘late’). For immune and glucocorticoid data, data were only collected from cold-induced dormancy organisms (*i.e.,* no available data for estivating organisms).

Overall, our dataset includes 2,840 unique estimates, from 112 unique papers (Kastner *et al*., 1978; Gustafson and Belt, 1981; Shivatcheva, Ankov and Hadji∼uff, 1987; Dauphin-Villemant, Leboulenger and Vaudry, 1990; Cooper *et al*., 1992, 2016; Hermes-Lima and Storey, 1995; Pérez-Pinzón and Rice, 1995; Grundy and Storey, 1998; Drew *et al*., 1999; Bitting *et al*., 1999; Wróbel, Sura and Srebro, 2000; Carey, Frank and Aw, 2000; Carey, Frank and Seifert, 2000; Chauhan *et al*., 2002; Maniero, 2002, 2005; Donahue *et al*., 2003; Ferreira, Alencastro and Hermes-Lima, 2003; Bagnyukova, Storey and Lushchak, 2003; Lushchak *et al*., 2005; Ma *et al*., 2005; Ramos-Vasconcelos, Cardoso and Hermes-Lima, 2005; Hudson *et al*., 2006; Ohta *et al*., 2006; Okamoto *et al*., 2006; Osborne and Hashimoto, 2006, 2007; Kurtz and Carey, 2007; Morin and Storey, 2007; Serkova *et al*., 2007; Fedotcheva *et al*., 2008; Lutterschmidt and Mason, 2008, 2009; Pier *et al*., 2008; Astaeva and Klichkhanov, 2009; Giroud *et al*., 2009, 2021; Nowakowska, Grazyna Świderska-Kołacz, *et al*., 2009; Nowakowska, Grázyna Świderska-Kołacz, *et al*., 2009; Orr *et al*., 2009; Page *et al*., 2009, 2010; Nowakowska, Caputa and Rogalska, 2010, 2011; Salway, Tattersall and Stuart, 2010; Giraud-Billoud *et al*., 2011, 2013, 2018, 2022; Bouma *et al*., 2011; Schwanz *et al*., 2011; Bouma *et al*., 2013; Brown *et al*., 2012; Talaei *et al*., 2012; Weitten *et al*., 2013; Xu *et al*., 2013, 2021; Young, Cramp and Franklin, 2013; Costanzo and Lee, 2013; Feidantsis, Anestis and Michaelidis, 2013; Franco, Contreras and Nespolo, 2013; James *et al*., 2013; Klanian, 2013; Moore *et al*., 2013; Sahdo *et al*., 2013; Afifi and Alkaladi, 2014; Avci *et al*., 2014; Nowakowska *et al*., 2014; Bohr, Brooks and Kurtz, 2014; Rouble, Tessier and Storey, 2014; Wu *et al*., 2015; Tessier *et al*., 2015; Yin *et al*., 2016; Liu *et al*., 2016; Bhunia *et al*., 2016; Nowakowska, Rogalska and Caputa, 2016; Samanta and Paital, 2016; Zhang, C. juan Niu, *et al*., 2017; Zhang, C. Niu, *et al*., 2017; Logan and Storey, 2018, 2021; Moreira *et al*., 2018, 2020, 2021; Niu *et al*., 2018; Sprenger, Tanumihardjo and Kurtz, 2018; Wei *et al*., 2018, 2019; Zena *et al*., 2019; Chazarin *et al*., 2019; Chen *et al*., 2019; Fan *et al*., 2019; Logan *et al*., 2020; Sahoo and Patnaik, 2020; Shi *et al*., 2020; Vella *et al*., 2020; Anegawa *et al*., 2021; Huber *et al*., 2021; Patnaik and Sahoo, 2021; Tang, Chen and Niu, 2021; Tessier, Breedon and Storey, 2021; Wang *et al*., 2021; Xie, Ahmad, Zuo, Wang, *et al*., 2022; Xie, Ahmad, Zuo, Xiao, *et al*., 2022; Frøbert *et al*., 2022; Lutterschmidt, Lucas and Summers, 2022; Miao *et al*., 2022; Mikes *et al*., 2022; Tsiouris and Flory, 2023; Dori *et al*., 2024; Vicente-Ferreira *et al*., 2024). In hibernating animals, we obtained 1,435 estimates for oxidative stress (845 and 590 estimates across dormancy and arousal states, respectively), 418 for immune parameters (224 and 194 estimates across dormancy and arousal states, respectively), 123 for glucocorticoids (95 and 28 estimates across dormancy and arousal states, respectively). In aestivating animals, we obtained 864 estimates for oxidative stress (477 and 387 estimates across dormancy and arousal states, respectively), and no data were available for immune parameters and glucocorticoids.

#### Meta-analytic effect sizes

We compared the physiological values of individuals either during the dormant or arousal states against their physiological values in euthermia. For each comparison, we calculated the standardized mean difference with heteroscedastic population variances (hereafter ‘SMD’; Bonett, 2008, 2009). We calculated SMD along with their associated sampling variances using the R function ‘*escalc*’ in the ‘metafor’ R package (v4.8.0; Viechtbauer, 2010). SMD was calculated so that positive values meant higher estimates during dormancy or arousal states compared to their values in euthermia, and *vice versa*.

#### Meta-analysis and meta-regressions

We handled the datasets, ran all analyses and produced visualisations using R (v.4.4.2; R Core Team, 2024). To evaluate the effect of cold- or heat-induced dormancy on physiological parameters, we fitted multilevel (intercept-only) meta-analyses for each response term (i.e., one model for each oxidative stress, immunity and glucocorticoid datasets). All meta-analytic models estimated five random intercept effects, publication identity (i.e., among-study variation), class, family, species (i.e., to account for taxonomic variation) and an observation ID term. For the intercept-only models, we estimated total heterogeneity (*I*^2^) following (Naganawa and Santos, 2012; Senior *et al*., 2016) as implemented in the R function ‘*i2*_(‘orchaRd’ R package v2.0; Nakagawa *et al*., 2021). Then, we fitted individual models to investigate the differences across thermoregulatory modes (endotherms *versus* ectotherms). In the case of oxidative stress and immune parameters, we also ran meta-regressions including physiological subcategories as moderators. Additionally, to investigate if physiological subcategories responded similarly between endotherms and ectotherms, we run meta-regression including physiological subcategory as a moderator in meta-regression including endotherm or ectotherm data respectively.

#### Publication bias

We evaluated evidence for small-study effects and time-lag effects, following (Nakagawa *et al*., 2022). For that, we ran additional uni-moderator multilevel meta-analytic models, one for each effect size investigated. Each of these models included as a single moderator either the square-root of the inverse of the effective sample size or the mean-centered year of study publication (Trikalinos and Ioannidis, 2005; Nakagawa *et al*., 2022). Variation explained by these moderators (i.e., R^2^_marginal_) was calculated using the R function ‘r*2_ml’* (‘orchaRd’ R package v2.0; Nakagawa *et al*., 2021).

## Results

### Effect of dormant and arousal states on redox status in cold-induced dormant organisms

A global model including all oxidative stress estimates across species during the dormant state, showed significant increases in the expression of regulatory pathways ([95% confidence interval; 95% CI hereafter] = 0.300 [0.050 – 1.233]; Figure S1), non-significant decreases of ROS (estimate [95% CI] = 0.300 [-1.122 – 0.055]; Figure S1), and no changes in oxidative damage and antioxidant responses (Figure S5A). However, such redox changes were differentially affected by thermoregulatory capacity. In endotherms, the dormant state resulted in significant reductions of oxidative damage (estimate [95% CI] = -0.464 [-0.924 – -0.005]; Figure 1A) and increases in the expression of regulatory mechanisms (estimate [95% CI] = 0.602 [0.04 – 1.164]; Figure 1A), as well as it included non-significant decreases in ROS production (estimate [95% CI] = -0.538 [-1.098 – 0.022]; Figure 1A), and no changes in antioxidant levels (Figure 1A). In ectotherms, the dormant state resulted in significant increases in oxidative damage (estimate [95% CI] = 0.552 [0.019 – 1.085]; Figure 1B) and no changes in the levels of antioxidants (Figure 1B).

**Figure 1.**
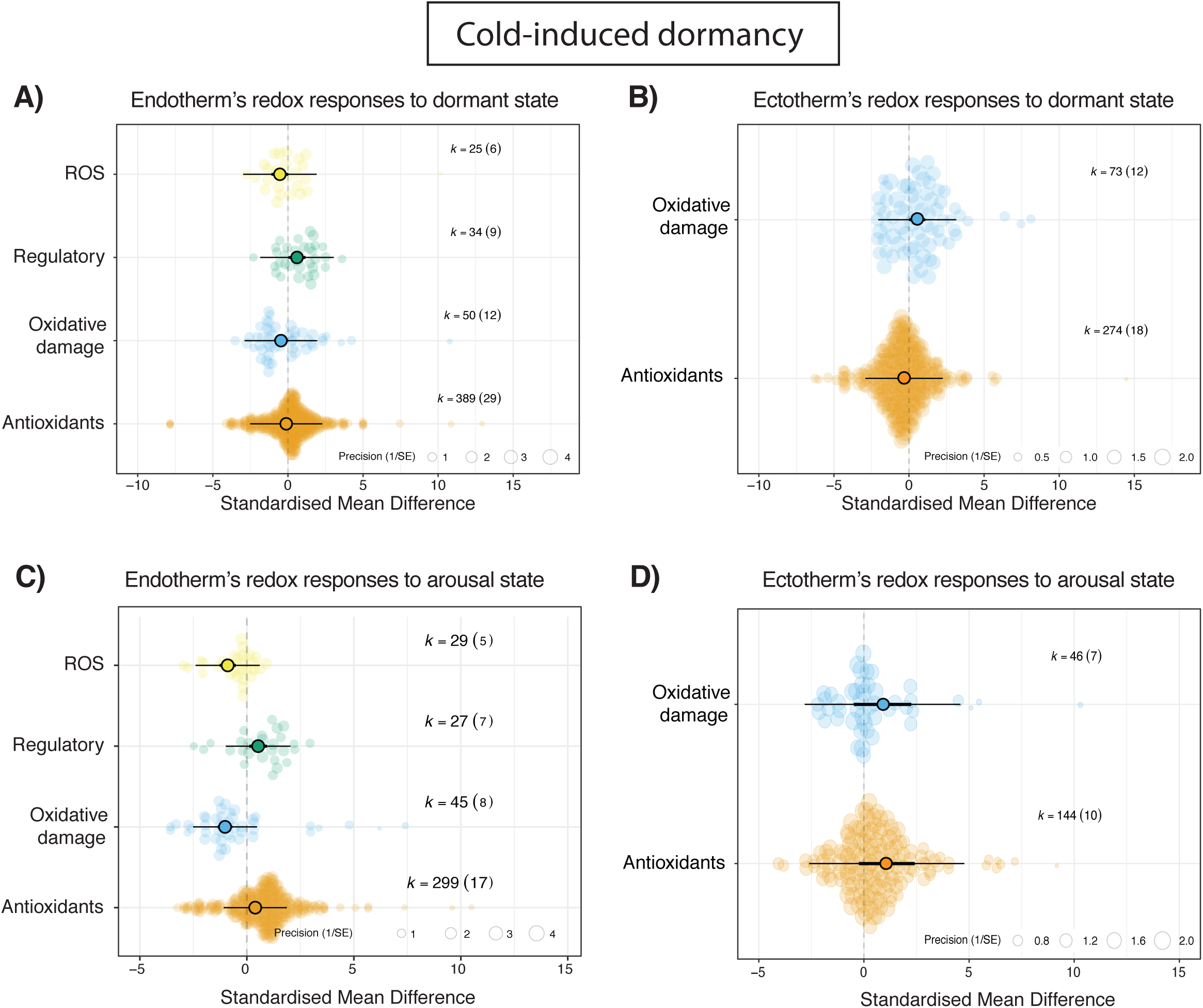
Orchard plots showing variation in redox status of endothermic and ectothermic animals to the dormant state (A and B, respectively) and arousal state (C and D, respectively) during cold-induced dormancy. We calculated variation in levels of reactive oxygen species (ROS), pathways involved the regulation of the redox balance, markers of oxidative damage (in lipids and proteins), and antioxidant defenses (enzymatic and non-enzymatic). Positive values on the X-axis indicate a higher value of a given component as compared to the euthermia state. Large coloured circles show overall means, a thick whisker 95% confidence interval, and a thin whisker 95% precision interval. The precision (1/SE) of each study is represented in the background of each panel, together with number of *k* estimates and studies (shown in parentheses).

During the arousal state, a global model reported an overall increase in the expression of regulatory pathways (estimate [95% CI] = 1.256 [0.062 – 2.450]; Figure S5B) and antioxidants (estimate [95% CI] = 1.074 [0.0.008 – 2.141]; Figure S1B), and no significant variation in ROS production or oxidative damage (Figure S5B). These patterns differed across parameters and animals regarding their thermoregulation capacity. During arousal, endotherms experienced significantly decreases in ROS production and oxidative damage (estimate [95% CI] = -0.893 [-1.391 – -0.395] and estimate [95% CI] = -1.018 [-1.466 – -0.395], respectively; Figure 1C) and increases antioxidant levels (estimate [95% CI] = 0.394 [0.078 – 0.709]; Figure 1C), and non-significant increases in the activity of regulatory pathways (estimate [95% CI] = 0.530 [-0.022 – 1.081]; Figure 1C). In contrast, ectotherms only experienced non-significant increases in markers of oxidative damage and antioxidants (Figure 1D).

### Effect of dormant and arousal states on redox status in heat-induced dormant organisms

During the dormant state, aestivating ectotherms experience significant increases in oxidative damage (estimate [95% CI] = 0.436 [0.077 – 0.795]; Figure 2) and no changes in ROS production, regulatory pathways, nor antioxidants (Figure 2A). In contrast, the arousal state did not influence redox parameters of aestivating animals (Figure 2B).

**Figure 2.**
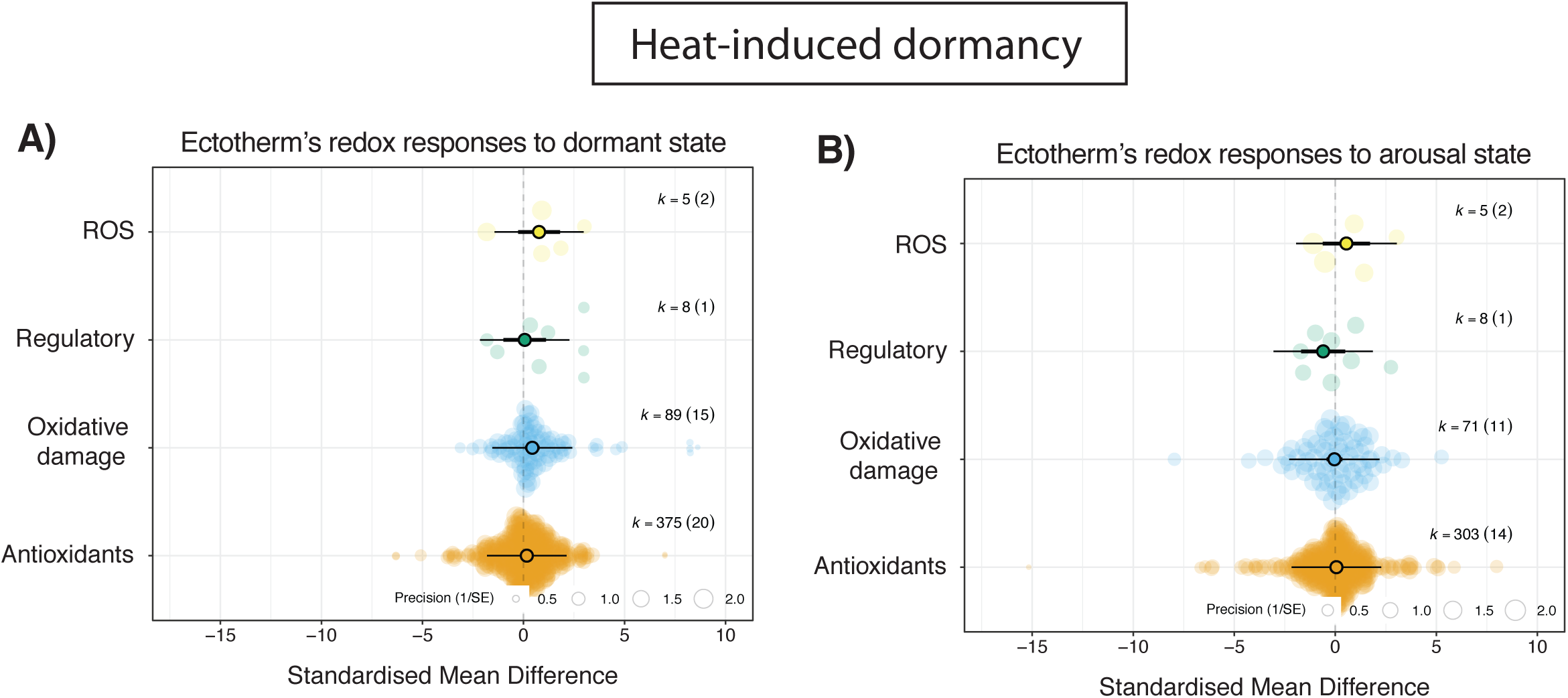
Orchard plots showing variation in redox status of ectothermic animals to the dormant state (A and B, respectively) and arousal state (C and D, respectively) during heat-induced dormancy (*i.e.,* aestivation). We calculated variation in levels of reactive oxygen species (ROS), pathways involved the regulation of the redox balance, markers of oxidative damage (in lipids and proteins), and antioxidant defenses (enzymatic and non-enzymatic). Positive values on the X-axis indicate a higher value of a given component as compared to the euthermia state. Large coloured circles show overall means, a thick whisker 95% confidence interval, and a thin whisker 95% precision interval. The precision (1/SE) of each study is represented in the background of each panel, together with number of *k* estimates and studies (shown in parentheses).

### Effect of dormant and arousal states on immune function in cold-induced dormant organisms

During the dormant state, a global model showed no significant variation in the different components of the immune system (*i.e.,* innate, adaptive, and both innate and adaptive responses; Figure S6A). However, the dormancy state had different effects on the immune system of endotherms and ectotherms. While in dormant endotherms no changes in any immune component were detected during (Figure 3A), ectotherms experienced significant downregulation of pathways involved in both innate and adaptive responses (*i.e.,* marginally non-significant response; estimate [95% CI] = -1.504 [-3.044 – 0.036]; Figure 3B), whereas it did not affect the innate immune function (Figure 3B).

**Figure 3.**
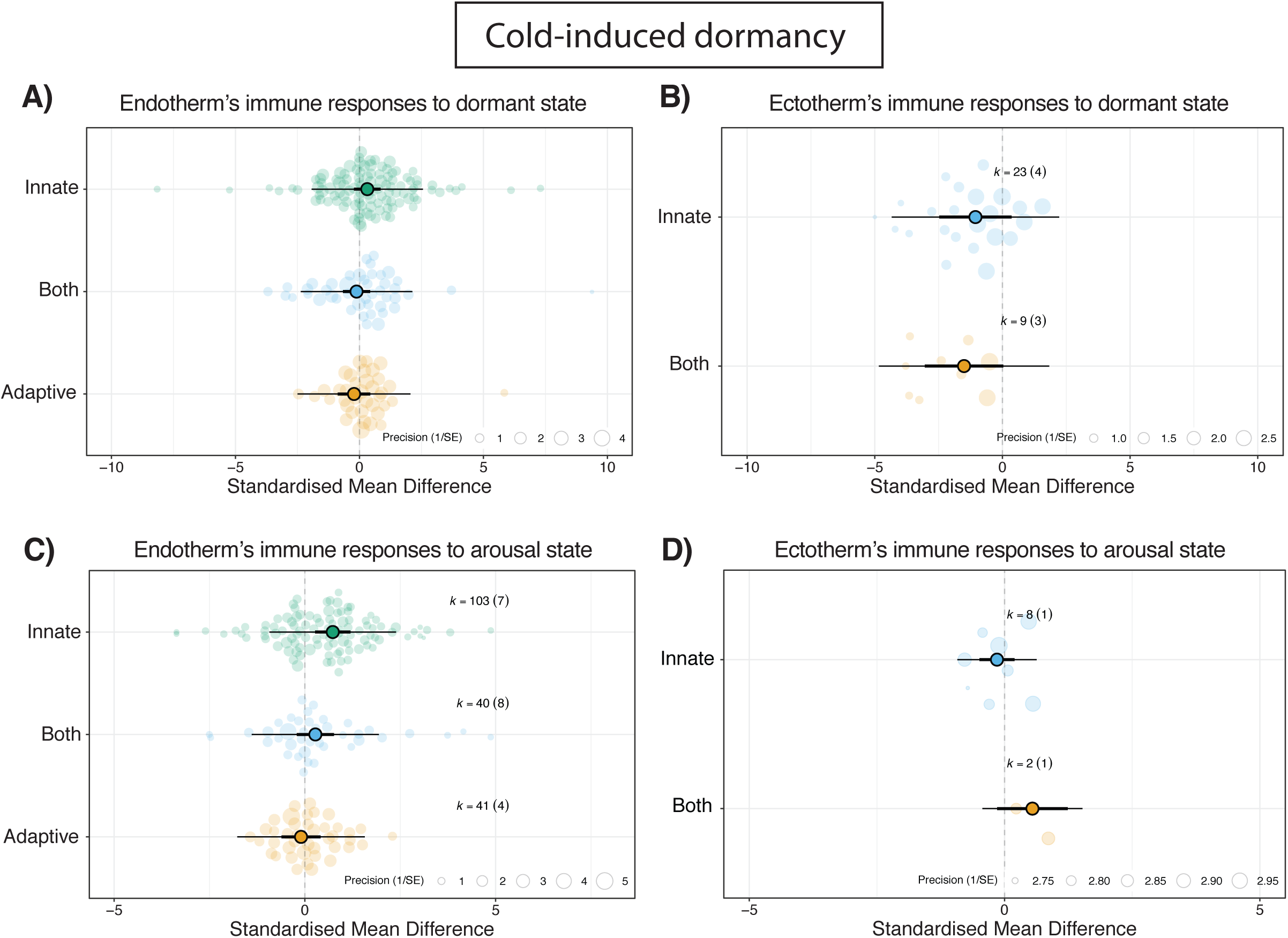
Orchard plots showing immune responses of endothermic and ectothermic animals to the dormant state (A and B, respectively) and arousal state (C and D, respectively) during cold-induced dormancy. We calculated variation on pathways involved either in the innate and adaptive immune responses, as well those considered as part of both the innate and adaptive immune response. Positive values on the X-axis indicate a higher value of a given component as compared to the euthermia state. Large coloured circles show overall means, a thick whisker 95% confidence interval, and a thin whisker 95% precision interval. The precision (1/SE) of each study is represented in the background of each panel, together with number of *k* estimates and studies (shown in parentheses).

A global model showed significant overall increases in the innate immune response during the arousal state (estimate [95% CI] = 0.577 [0.156 – 0.997]; Figure S6B). In endotherms, arousal resulted in upregulated innate immune response (estimate [95% CI] = 0.731 [0.268 – 1.194]; Figure 3C) and no significant variation was observed in the other immune components (Figure 3D). Data on immune responses in ectotherms undergoing arousal could not be meta-analysed since only one study met our criteria.

### Effect of dormant and arousal states on glucocorticoids in cold-induced dormant organisms

A global model showed no effect of the dormant state on glucocorticoid levels (estimate [95% CI] = 0.427 [-0.922 – 1.777]; Figure S6A), regardless of the thermoregulatory strategy of species (Figure 4A, 4B). Likewise, we did not find an overall effect of the arousal state on glucocorticoid levels (estimate [95% CI] = 0.427 [-0.922 – 1.777]; Figure S6B), nor in endotherms or ectotherms (Figure 4A, 4B).

**Figure 4.**
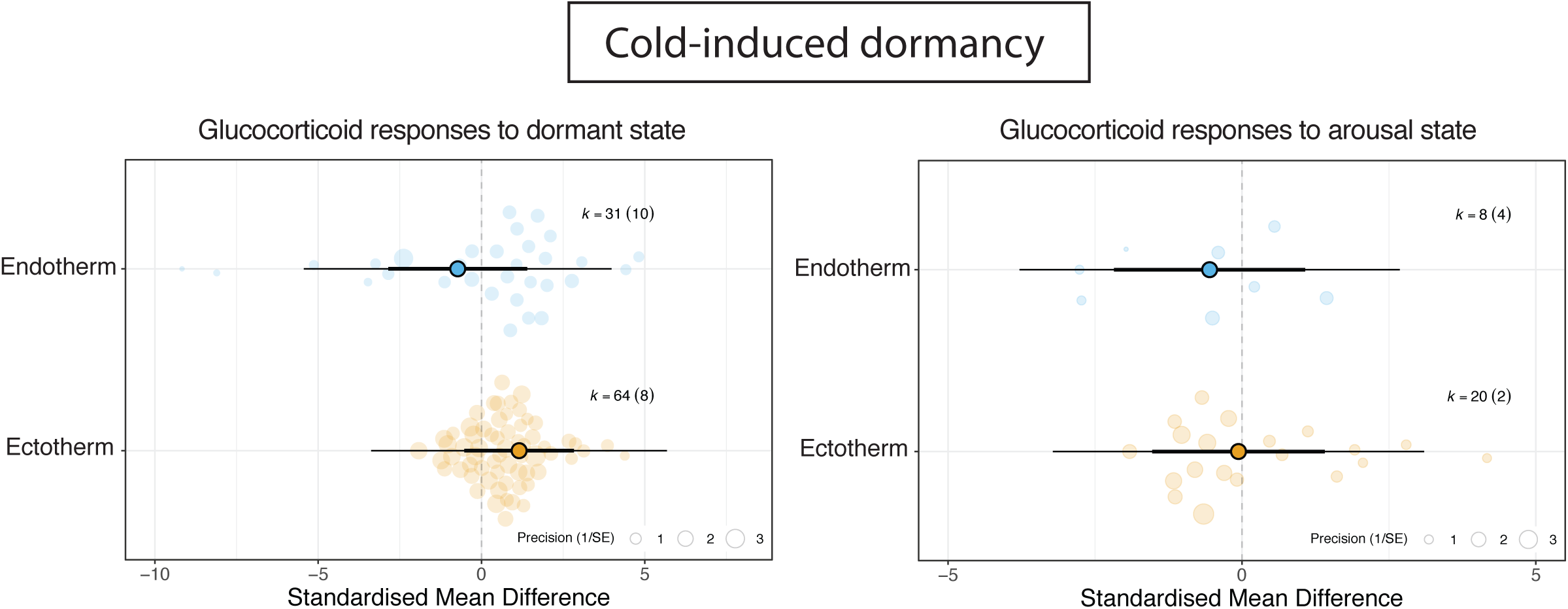
Orchard plots showing changes in glucocorticoid levels (cortisol and corticosterone) in endothermic and ectothermic vertebrates undergoing either the dormant state (A and B, respectively) and arousal state (C and D, respectively) cold-induced dormancy. Positive values on the X-axis indicate a higher value of a given component as compared to the euthermia state. Large coloured circles show overall means, a thick whisker 95% confidence interval, and a thin whisker 95% precision interval. The precision (1/SE) of each study is represented in the background of each panel, together with number of *k* estimates and studies (shown in parentheses).

## Discussion

Our study reveals contrasting effects of the dormancy states on animal redox status and immune function, which vary markedly depending on the organisms’ thermoregulatory strategy. In ectotherms, the dormant state is associated with a lowered redox status evidenced by higher oxidative damage, both during cold- and heat-induced dormancy. In contrast, endotherms experience enhanced redox status during the dormant and arousal states, indicated by reduced ROS production and oxidative damage, and higher antioxidant capacity. Regarding immune function, the dormant state induces subtle downregulation of pathways involved in both innate and adaptive responses in ectotherms, whereas arousal resulted in upregulated innate responses in endotherms. Finally, glucocorticoid release was not influenced by the dormancy process. Overall, dormancy remarkably influences animal physiology, with endotherms showing signs of enhanced physiology and ectotherms being negatively affected.

Metabolic shifts that occur during dormancy are often thought to abruptly alter energy allocation and lead to physiological stress (Ali *et al*., 2010; Brown *et al*., 2012; Staples, Mathers and Duffy, 2022). Our meta-analysis partially supports this idea, with responses being different according to thermoregulatory strategy of organisms. This is especially evident regarding changes in levels of markers of oxidative stress and damage. Endotherms experience decreases in ROS production and oxidative damage as well as upregulations in antioxidant capacity and expression of regulatory pathways involved in the protection from oxidative damage during the dormancy and arousal states. In contrast, ectotherms experience oxidative damage during the dormancy state (a pattern observed both in cold- and heat-induced dormant organisms), and negligible variation in their antioxidant system during the arousal state. Overall, changes in the redox status seem to reflect fundamental differences in how endotherms and ectotherms have evolved to cope with metabolic shifts during dormancy. The capacity of endotherms to actively regulate their internal temperature may allow for a more controlled depression and reactivation of metabolism (Clarke and Pörtner, 2010), having beneficial effects on their redox status which is also favoured by upregulated antioxidant pathways. These findings may mechanistically explain longer lifespans observed in dormant endotherms as compared to non-dormant endotherms (Turbill, Bieber and Ruf, 2011; Wu and Storey, 2016). The fact that ectotherms’ body temperature is largely dictated by environmental temperature could make them to enter dormancy more passively and with less metabolic control, finally resulting in an oxidative stress state. Costs associated to dormancy could impose strong selective pressures on ectothermic animals, potentially impacting on ectotherm health and lifespan, as well as influencing the duration and timing of the dormancy process and other aspects such as habitat selection. Indeed, oxidative stress is often presented as a main mediator of life-history trade-offs, including phenotypic responses to environmental shifts (Monaghan, Metcalfe and Torres, 2009; Metcalfe and Alonso-Alvarez, 2010; Burraco *et al*., 2022). Together, our findings highlight how thermoregulatory strategy influences the physiological consequences of dormancy which may explain life-history evolution, but the latter needs further exploration.

Oxidative stress can contribute to immune dysfunction, leading to chronic inflammation with potentially serious consequences for health and fitness (Sorci and Faivre, 2009; Lauridsen, 2019; Costantini, 2024). In our study, consistently with changes in endotherms oxidative status, we found that endothermic organisms experience from negligible changes in immune function during the dormancy state to upregulated innate immune responses during arousal. These findings further support that dormancy process has few physiological costs (if not beneficial) for endothermic organisms. However, such conclusion must be made cautiously, given the complexity of immune response and their context-dependent regulation. Indeed, research has demonstrated dramatic consequences for some hibernating endothermic species. For instance, the white-nose syndrome, a fungal infection, is known to devastate hibernating bat populations by causing irritation on bats that prompts more frequent arousals, leading to premature depletion of energy stores (Reeder *et al*., 2012). In dormant ectotherms, also consistently with observed increased oxidative stress, our meta-analysis shows that the dormancy state downregulates the expression of pathways involved in both the innate and adaptive immune response. This likely reflects a physiological trade-off, in which energy and resources are reallocated away from costly immune functions to sustain other essential functions under metabolically suppressed conditions (Pörtner *et al*., 2006; Butler *et al*., 2013; Barley *et al*., 2021). Such immunosuppression may result in a higher vulnerability to infections and reductions in their ability to respond effectively to pathogens upon arousal. However, this idea needs further evidence since only few studied have been so far conducted in this direction. Overall, our meta-analytic approach further reinforces the idea that dormancy impairs the physiology of ectotherms whereas impacts on endotherms are minor.

Glucocorticoid release is known to be key regulator of metabolism or energy mobilisation, and can to physiological alterations via changes in the production of ROS or antioxidant molecules (*e.g.* glutathione), or immune function (*e.g.,* by inhibiting pro-inflammatory cytokines or altering the activity of immune cells such as T cells) (Bauer, 2005; Costantini, Marasco and Møller, 2011; Angelier and Wingfield, 2012). Given the link between glucocorticoids and metabolism, dormancy-related transitions are expected to govern shifts in glucocorticoids release. Our meta-analysis shows that glucocorticoid levels remain stable across dormancy and arousal states, even though redox balance and immune status undergo marked changes in both endotherms and ectotherms. This may suggest that dormant species could have evolved the ability to decouple metabolism from glucocorticoid release (Vitousek *et al*., 2019), thereby buffering the cascade of physiological consequences. Alternatively, time-dependent or tissue-specific fluctuations may have not been captured in the available data (Crespi *et al*., 2013; Romero and Beattie, 2022). Yet, knowledge on glucocorticoid dynamics across dormancy states is still restricted to some species and should be complemented with further research.

While our meta-analysis provides a broad overview of how dormancy phases affect physiological parameters across taxa, several important questions remain unresolved due to data limitations. For instance, current evidence does not yet allow us to disentangle how different stages within the dormancy process (e.g., early vs. late) influence physiological responses, or how the number and frequency of dormancy events, their interaction with other stressors, or variation across life stages (e.g., Martin *et al*., 2023) and life-history traits (e.g., sex, body size) may modulate these effects. In addition, key physiological indicators such as telomere dynamics remain largely unexplored in this context, despite recent reviews highlighting their potential relevance (Redon *et al*., 2024). Addressing these gaps will be crucial to refine our understanding of the impacts of dormancy in wild organisms and to enhance future assessments and predictions of population status, including through the use of mechanistic species distribution models (SDMs) (Kearney and Porter, 2009; Talluto *et al*., 2016; Becker *et al*., 2020; Burraco, Lucas and Salmón, 2022).

## Conclusions

Our meta-analysis shows that dormancy includes contrasting physiological effects in dormant animals with different thermoregulatory strategies. While dormant endotherms overall experience enhanced physiological status, the dormancy process negatively impacts on ectotherms’ physiology. This knowledge aims to improve our mechanistic understanding of the influence of dormancy on animal condition and health, which aims to improve future research on eco-evolutionary processes in dormant species. Also, our meta-analysis may have potential applications in research on metabolic depression, and inform conservation plans.

## SUPPLEMENTARY MATERIAL

**Figure S1.**
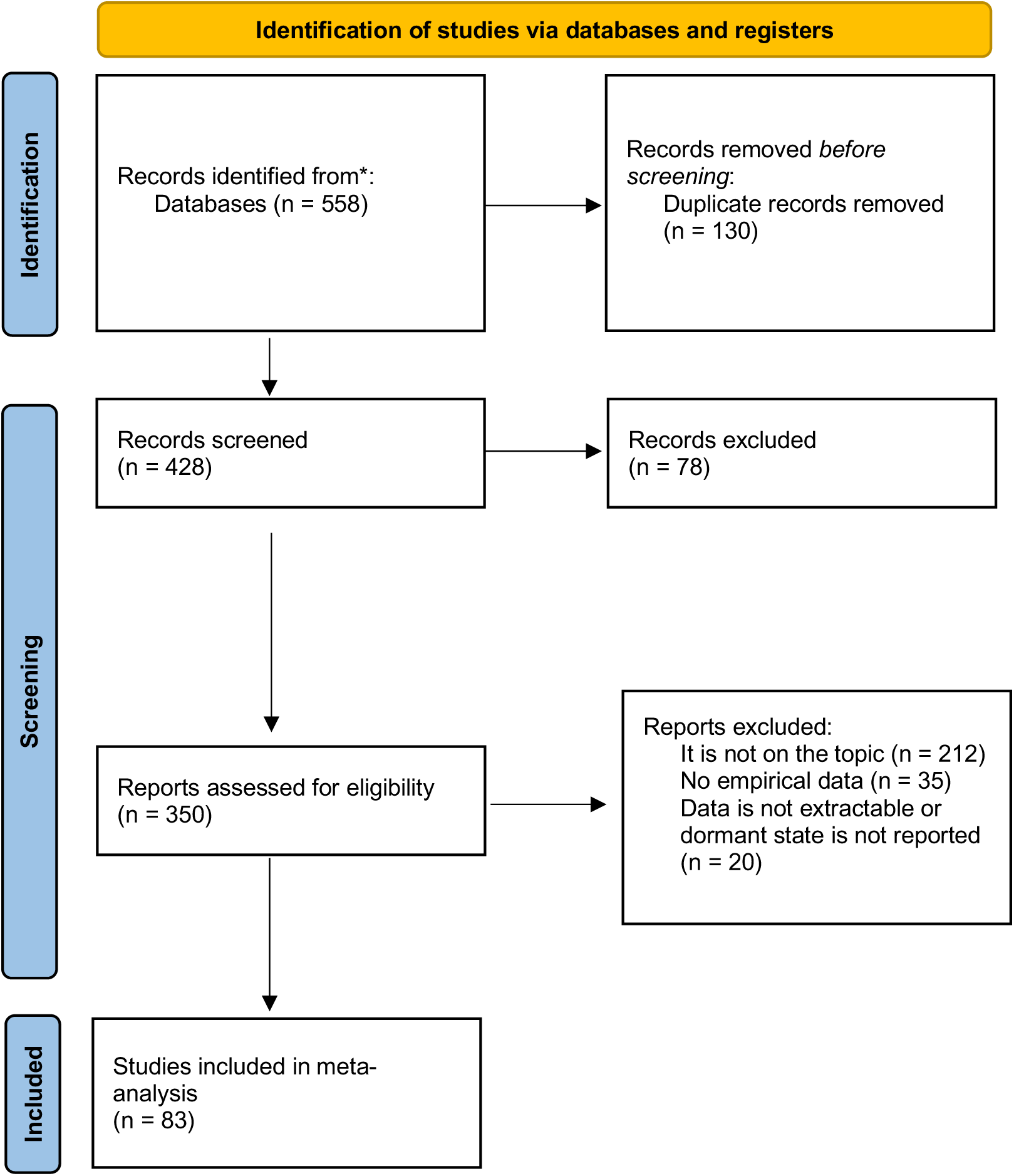
PRISMA flow diagram illustrating the study selection process for the meta-analysis assessing the effects of cold- and heat-induced dormancy on oxidative stress markers. The diagram summarizes the number of records identified through database searching (*i.e.,* Web of Science, Scopus, Pubmed), screened, assessed for eligibility, and ultimately included in the quantitative synthesis.

**Figure S2.**
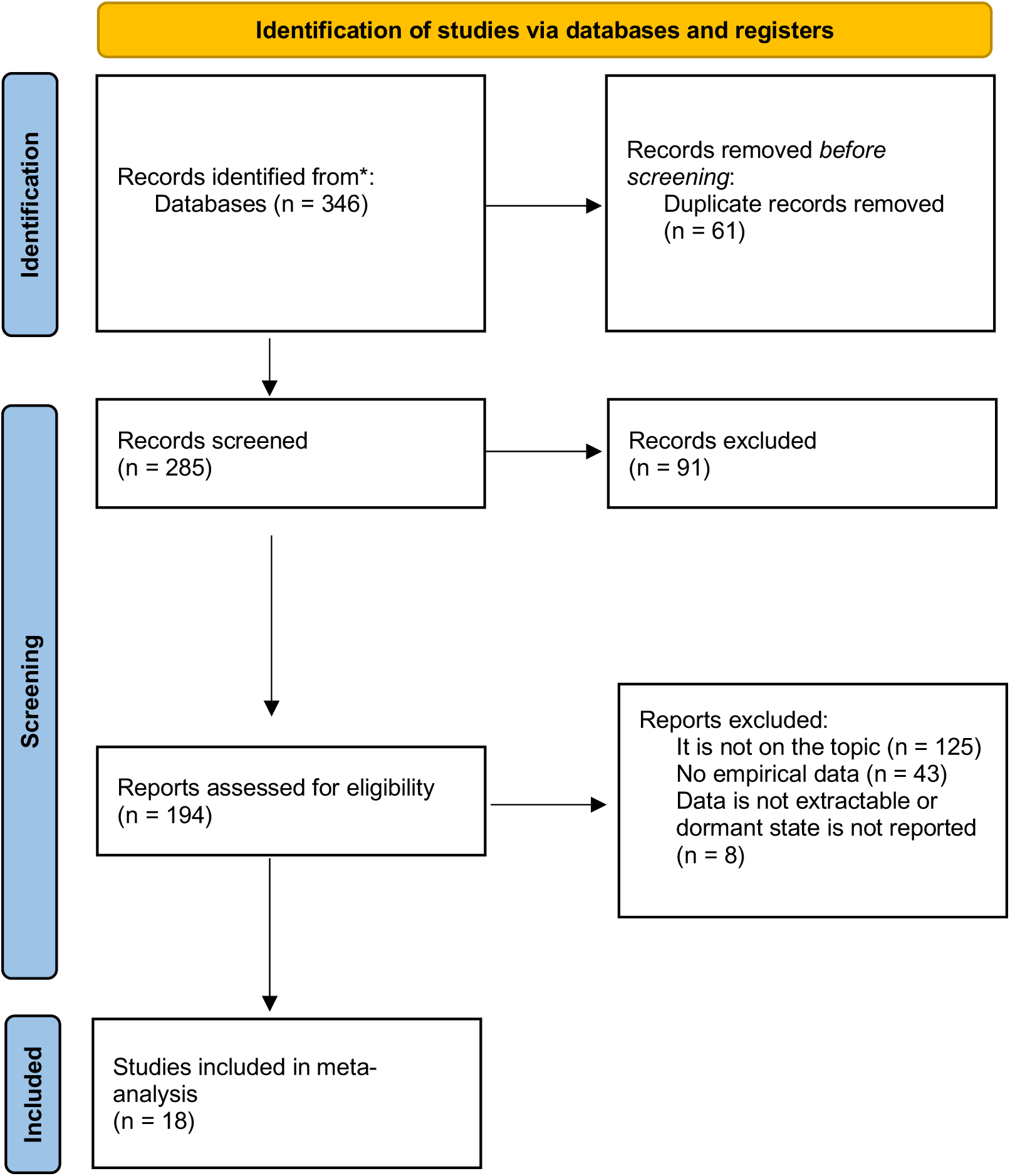
PRISMA flow diagram illustrating the study selection process for the meta-analysis assessing the effects of cold- and heat-induced dormancy on immune parameters. The diagram summarizes the number of records identified through database searching (*i.e.,* Web of Science, Scopus, Pubmed), screened, assessed for eligibility, and ultimately included in the quantitative synthesis.

**Figure S3.**
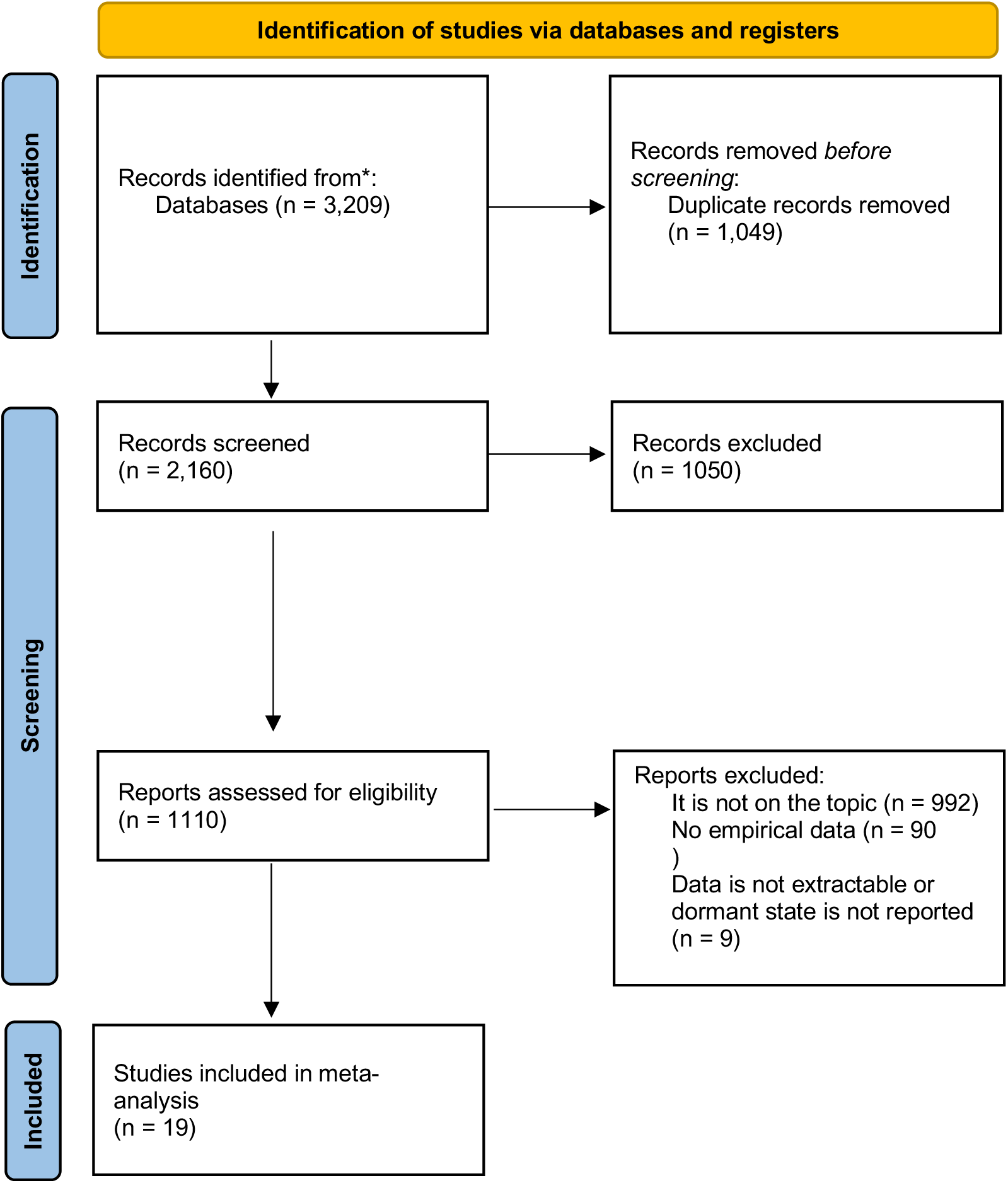
PRISMA flow diagram illustrating the study selection process for the meta-analysis assessing the effects of hibernation, torpor, or aestivation on glucocorticoids. The diagram summarizes the number of records identified through database searching (*i.e.,* Web of Science, Scopus, Pubmed), screened, assessed for eligibility, and ultimately included in the quantitative synthesis.

**Figure S4.**
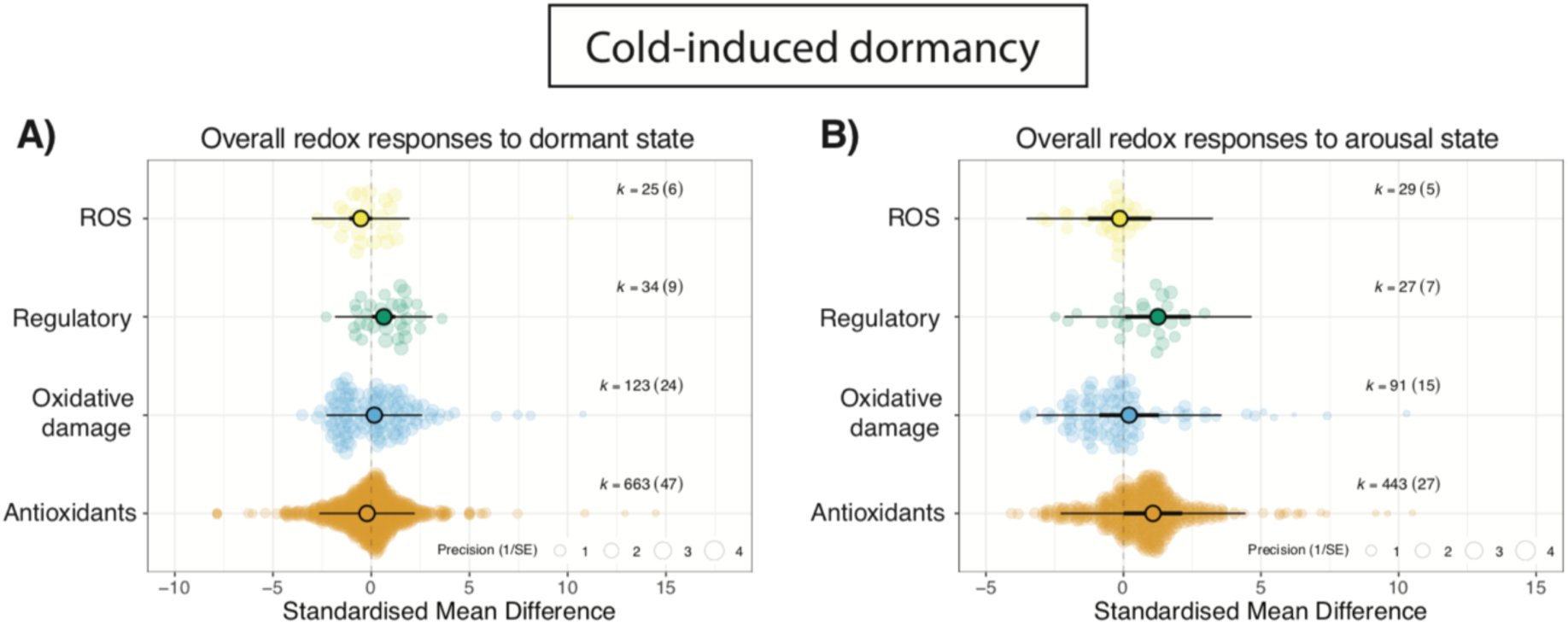
Orchard plots showing overall variation in redox status in animals (*i.e.,* both endotherms and ectotherms) undergoing A) the dormant state (A and B, respectively) and B) the arousal state during cold-induced dormancy. We calculated variation in levels of reactive oxygen species (ROS), pathways involved the regulation of the redox balance, markers of oxidative damage (in lipids and proteins), and antioxidant defenses (enzymatic and non-enzymatic). Positive values on the X-axis indicate a higher value of a given component as compared to the euthermia state. Large coloured circles show overall means, a thick whisker 95% confidence interval, and a thin whisker 95% precision interval. The precision (1/SE) of each study is represented in the background of each panel, together with number of *k* estimates and studies (shown in parentheses).

**Figure S5.**
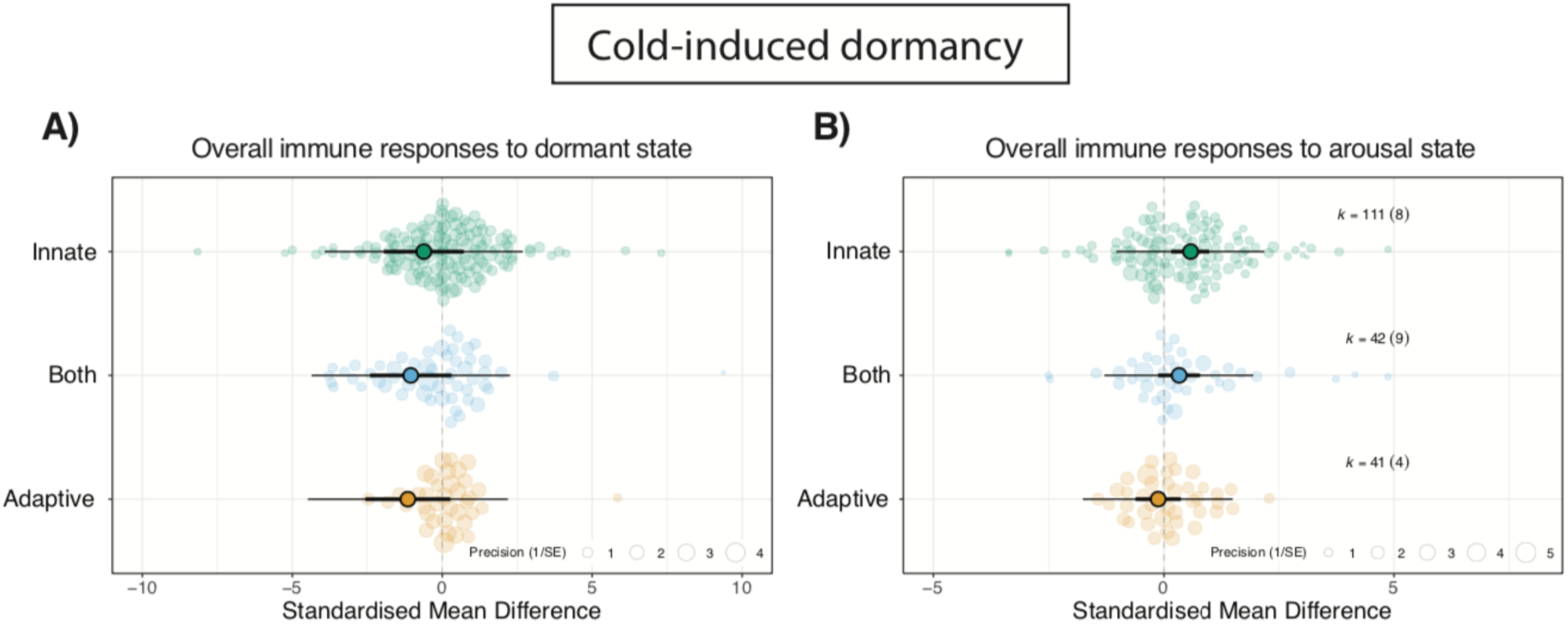
Orchard plots showing immune responses in animals (*i.e.,* both endotherms and ectotherms) undergoing A) the dormant state (A and B, respectively) and B) the arousal state during cold-induced dormancy. We calculated variation on pathways involved either in the innate and adaptive immune responses, as well those considered as part of both the innate and adaptive immune response. Positive values on the X-axis indicate a higher value of a given component as compared to the euthermia state. Large coloured circles show overall means, a thick whisker 95% confidence interval, and a thin whisker 95% precision interval. The precision (1/SE) of each study is represented in the background of each panel, together with number of *k* estimates and studies (shown in parentheses).

**Figure S6.**
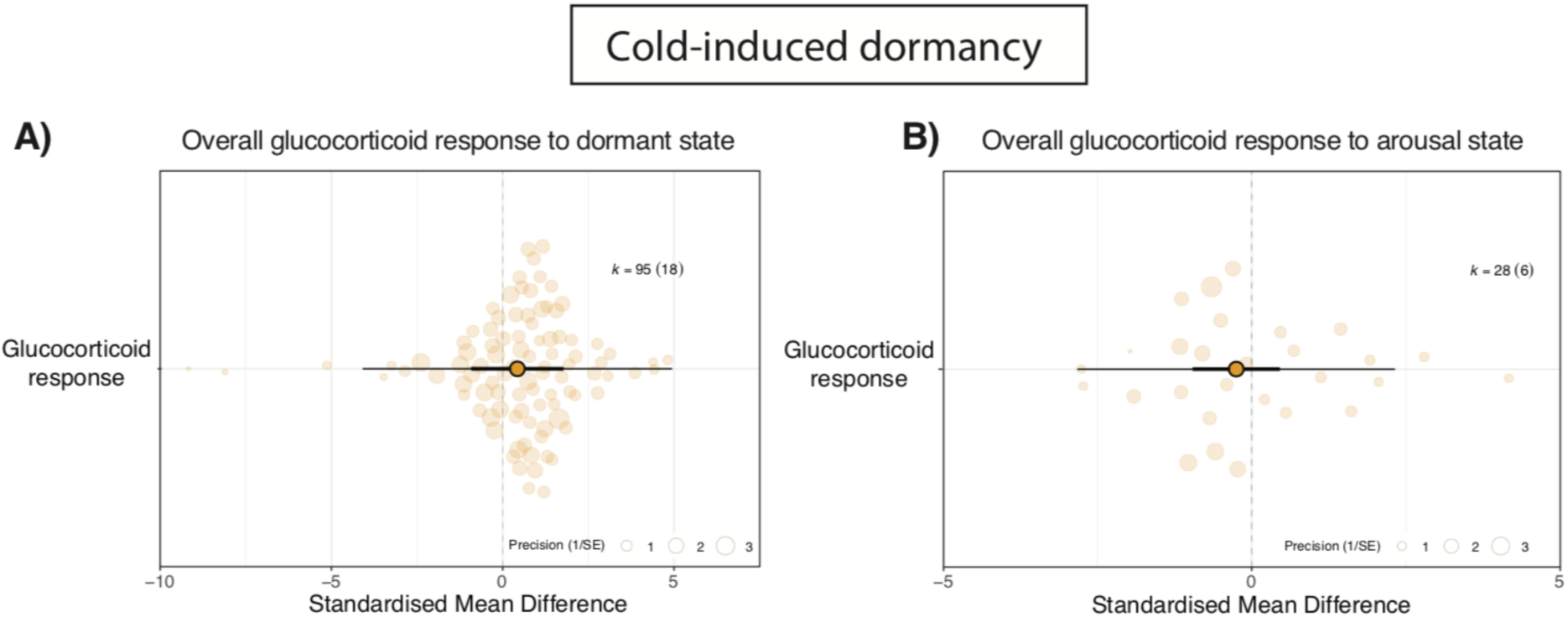
Orchard plots showing changes in glucocorticoid levels (cortisol and corticosterone) in in animals (*i.e.,* both endotherms and ectotherms) undergoing A) the dormant state (A and B, respectively) and B) the arousal state during cold-induced dormancy. Positive values on the X-axis indicate a higher value of a given component as compared to the euthermia state. Large coloured circles show overall means, a thick whisker 95% confidence interval, and a thin whisker 95% precision interval. The precision (1/SE) of each study is represented in the background of each panel, together with number of *k* estimates and studies (shown in parentheses).

## Notes

### Competing Interest Statement

The authors have declared no competing interest.

